# Orally delivered biodegradable targeted inflammation resolving pectin-coated nanoparticles induce anastomotic healing post intestinal surgery

**DOI:** 10.1101/2024.01.01.569918

**Authors:** Jong Hyun Lee, Stefan Reischl, Robert Leon Walter, Vincent Vieregge, Marie-Christin Weber, Runxin Xu, Hao Chen, Atsuko Kasajima, Helmut Friess, Philipp-Alexander Neumann, Nazila Kamaly

**Author notes:** These authors contributed equally to this work (Philipp-Alexander Neumann, Nazila Kamaly). These authors contributed equally to this work (Jong Hyun Lee, Stefan Reischl).

## Abstract

Although medical treatment is sucessful in most cases in patients with inflammatory bowel diseases (IBD), a percentage of patients require surgical resection of diseased bowel segments at least once in their lifetime. Healing success of the intestinal anastomosis is at high risk, especially in presence of acute inflammation. Failure of anastomotic healing is a life-threatening complication and causes high socioeconomic costs. Common anti-inflammatory medications can have detrimental effects on wound healing. Thus, targeted perioperative therapeutics supporting anastomotic healing during colitis are an urgent medical need. Here, we develop a novel basal membrane targeted controlled release, pectin-coated polymeric nanoparticle (NP) encapsulating a highly potent inflammation resolving mediator, the peptide Ac2-26. These NPs can undergo gastric passage and facilitate localized release of the therapeutic peptide in the colon via degradation of their pectin-chitosan coating by microbial pectinases, which subsequently exposes a collagen IV targeted NP surface, allowing for further binding and retention of the NPs at the intestinal wound. To test these NPs, we used a murine surgical model combining the formation of an intestinal anastomosis with the induction of a preoperative colitis by dextran sodium sulfate. In this model, perioperative administration of pectin-chitosan coated NPs containing Ac2-26 (P-C-Col IV-Ac2-26-NP) led to the reduction of colitis activity in the postoperative phase. Macroscopic wound closure was improved by P-C-Col IV-Ac2-26-NP treatment as evaluated by endoscopy and intraabdominal adhesion scoring. Microscopic analysis of the healing process showed an improved semiquantitative healing score in the treatment group. In this proof-of-concept study we demonstrate that novel P-C-Col IV-Ac2-26-NP could be a promising and clinically feasible perioperative treatment strategy for IBD patients.

**TOC graphic:** 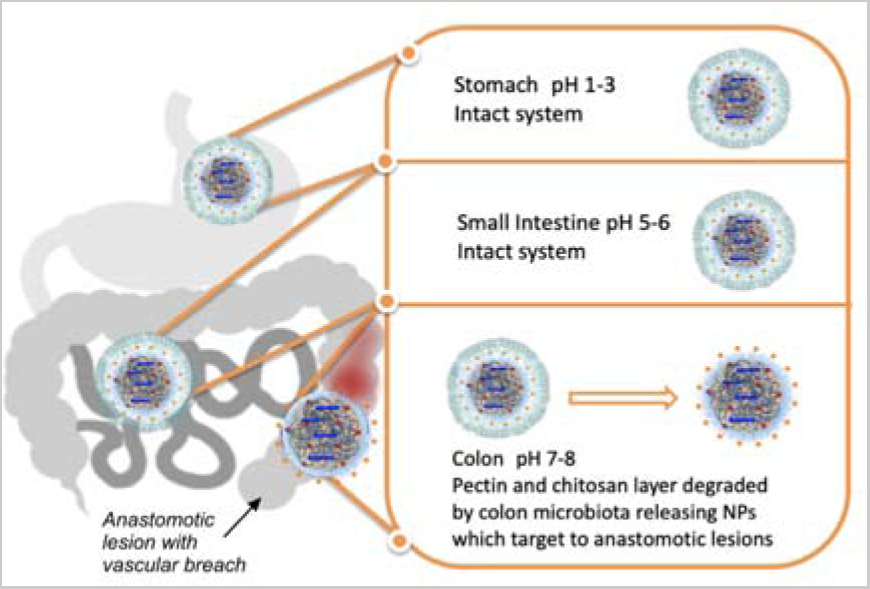

## INTRODUCTION

Inflammatory bowel diseases (IBD) - comprising ulcerative colitis and Crohńs disease - represent chronic inflammatory disorders of the gastrointestinal tract that are associated with relapsing symptoms of severe diarrhea, fever, weight loss, and abdominal pain.^1, 2^ Immunosuppressive medical therapy is considered the first line approach for treatment of IBD. Although medical therapy is successful in most IBD patients, over 40% require surgical interventions at least once in their lifetime.^1,2^ Surgical anastomoses to reconnect the bowel after resection of an intestinal segment are frequent procedures used in IBD surgery. Despite technical advances in surgery, postoperative anastomotic leakage occurs in up to 20% of colorectal operations and leads to high morbidity.^3^ Intestinal inflammation as seen in these patients can disturb the healing process, often leading to failed anastomotic healing.

The attempt to dampen the inflammation caused by active colitis prior to surgery by immunosuppressive drugs such as glucocorticoids or TNF-alpha inhibitors might even further disturb a regulated healing process after anastomosis formation. This is because throughout the anastomotic healing process, a precise balance of pro- and anti-inflammatory signaling is essential for successful healing. The initial inflammatory reaction seems to be indispensable, as treatment with non-steroidal anti-inflammatory drugs strongly reduces the inflammatory response, leading to an increase in anastomotic leakage rates.^4^

The resolution of inflammation is innately mediated by various lipids and proteins, which block inflammatory cell influx, promote their egress, clear pathogens, cellular debris and inflammatory cytokines, and repair tissue damage.^5, 6^ Resolution mediators include endogenous lipids that are generated during inflammation, e.g. lipoxins, resolvins, protectins, maresins; and proteins such as Annexin A1 (ANXA1), which is a key protein that is capable of facilitating resolution of inflammation.^7, 8^ This protein binds to the formyl peptide receptor expressed on activated cells such as phagocytes and epithelial cells and can resolve inflammation. It exerts inflammation resolution by suppression of pro-inflammatory signaling (e.g. NF-κB signaling).^9, 10^ It has shown pro-resolving properties in various inflammation models that include endotoxemia, peritonitis and arthritis.^11–13^ Furthermore, intraperitoneal injections of recombinant ANXA1 lead to closure of mucosal wounds induced with biopsy forceps and a reduction of colitis activity induced by dextran sodium sulfate.^14^ These results make ANXA1 and its pharmacophore, the 26 amino acid *N*-terminal peptide Ac2-26^15–17^ a promising candidate for perioperative treatment of IBD patients.

Although oral administration would be a cost-effective route to locally deliver biological therapeutics to influence the healing process of intestinal wounds, degradation during gastric passage complicates this strategy. Nanoparticle (NP) drug delivery systems allow encapsulation of the therapeutic agent, and can protect biological payloads during gastric passage if gut resistant compositions are used. Amongst the most successfully translated nanomedicines are drug loaded polymeric NPs, which have proven to be promising tools for the specific and sustained delivery of a range of therapeutics.^18–21^ Polymeric NPs are also currently being investigated for oral delivery applications.^24, 25^ NP-mediated delivery of biologics, including resolution mediators, can facilitate their: (a) targeted delivery; (b) controlled release, which facilitates dose tempering; and (c) protection from stomach components until release at target site. Furthermore, pectin-chitosan coating of polymeric NPs allows for protection against degradation during gastric passage with localized release occurring after specific pectin digestion in the colon by microbial pectinases and colonic bacteria.^26–28^ We have previously shown that intraperitoneal application of polymeric Ac2-26-NPs is effective in inducing intestinal wound healing processes and recovery from experimental colitis.^29–31^ These results suggest a high therapeutic potential for NP encapsulated Ac2-26 to induce anastomotic healing during colitis.

In this current work, we aimed to further develop these NPs for oral administration by coating them initially with a chitosan layer, followed by a subsequent pectin layer for degradation within the colon and release of the core collagen IV targerted Ac2-26 NPs (Figure 1A). The orally administered NPs have to withstand various environmental conditions in the gastrointestinal tract (GIT), a low pH in the stomach, mechanical pressure, protease attack in the small intestine and microflora digestion in the colon. In order to minimize the impact on the stability of the Ac2-26 peptide and the targeting collagen IV peptide on the surface of the NPs, following a chitosan coating, we further coated the NPs with pectin, which is a naturally occurring edible polysaccharide with a long and safe history of use in the food industry.^32^ In fact, pectin passes intact through the upper GI tract and is degraded by colonic microflora, which will ensure degradation and NP release at the anastomotic site within the colon. Pectin on its own cannot be used to deliver the naked Ac2-26 peptide, as pectin swells considerably in physiological conditions and the peptide will be released prematurely through the large pores. We therefore encapsulated polylactic-co-glycolic acid (PLGA) NPs entrapping the Ac2-26 peptides with a short PEGylated surface and collagen IV targeting peptides within chitosan and subsequent pectin layers and tested these in anastomosis models. The additional chitosan and pectin layers help protect the collagen IV targeting peptide ligand, in addition to the Ac2-26 peptide until the NPs have reached the lower colon where peptinases and colon microbiota can degrade their pectin and chitosan layers, releasing the Col IV-Ac2-26 NP to target the anasotomotic site (Figure 1B).

**Figure. 1.**
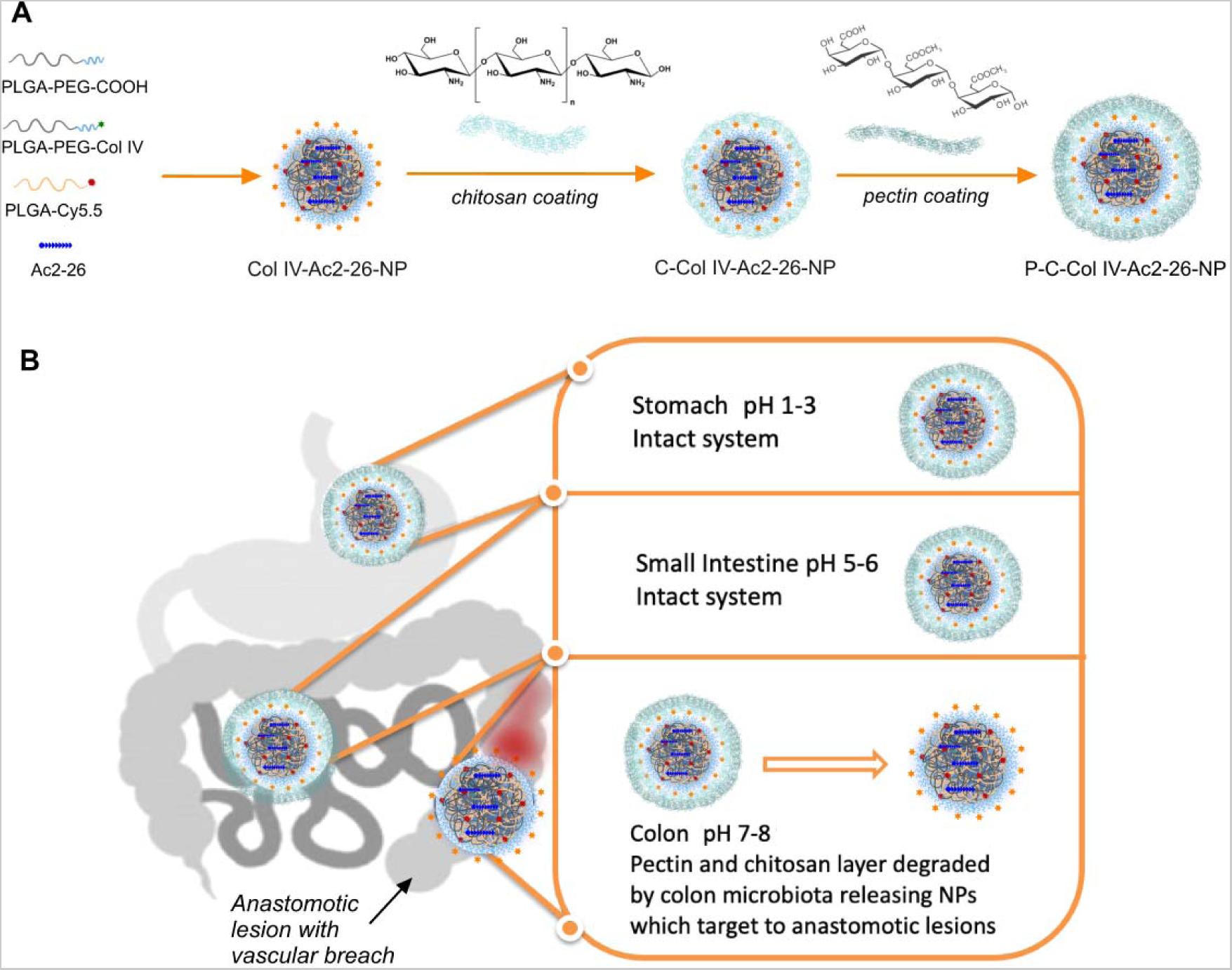
Overall design of biodegradable layer-by-layer P-C-Col IV Ac2-26 NPs for anastomotic healing. The pectin coating of the NPs protects from the harsh pH environment of the stomach, small intestine and enzymes secreted for digestion. Once in the colon, pectin and the subsequent chitosan layers are fermented and degraded almost completely by the microflora in the gut and this leads to the local release of the Col-IV Ac2-26 NPs, which binds to wounds and releases the Ac2-26 peptide *in situ*.

## RESULTS AND DISCUSSION

### Development of pectin-coated polymeric Ac2-26 loaded nanoparticles (P-C-Col IV-Ac2-26 NPs)

We have previously shown that collagen-IV-targeted Ac2-26 containing nanoparticles (Col IV-Ac2-26-NPs) are inflammation resolving in murine peritonitis, hind-limb reperfusion injury, advanced atherosclerosis, mucosal wound repair and experimental colitis.^10, 20, 29, 30^ In this study, we developed a modified version of these NPs with a chitosan and pectin coating for oral delivery and investigated their efficacy in our anastomotic healing model during colitis. Most importantly, the use of controlled release polymeric NPs can facilitate dose tempering of the potent pro-resolving Ac2-26 peptide, which suffers from short plasma-half life and non-specific effects when administered in its free form. In the first instance we synthesised the required copolymers of poly[lactic-*co*-glycolic acid]-*b*-poly[ethylene glycol] (PLGA-PEG) and PLGA-PEG-Col IV as previously reported.^30^ The major feature of these polymers is that they utilise a very short PEG sequence of about 534.6 Da in order to allow for chitosan coating, which provides the positive electrostatic interaction with the negatively charged pectin polymer – both chitosan and pectin are degradable once the NPs have reached the colon. For these NPs, a collagen IV (Col IV) targeting approach was used to retain the Ac2-26-NPs (post chitosan and pectin biodegradation) at inflamed or wounded intestinal tissue and to enable the release of the Ac2-26 peptide. Col IV is a major component of the basal membrane, and is exposed at sites of tissue injury, hence serving as an optimal extracellular matrix targeting strategy.^33^

In addition to Col IV-targeted Ac2-26 NPs, targeted NPs containing a randomly generated, isoelectric miss-matched scrambled peptide sequence to Ac2-26 (Scrm-NP) were also formulated as controls. The NP encapsulation efficiency for the Ac2-26 peptide and scrambled peptide were 87.5 and 91.7%, respectively. The loading efficiency for the Ac2-26 peptide and scrambled peptide were 10.5 and 11%, respectively. Col IV Ac2-26 NPs and Col IV Scrm NPs had a size distribution between 149.0 ± 2.5 nm and 155.25 ± 3.7 nm, indicating these two NPs are close in size for effective comparison (Figure 2A). We then observed an increase in size for these NPs following electrostatic coating with chitosan up to approximately 350 nm, indicating extra layers existing on the surface of the NPs following this intial coating (Figure 2A). Next, the NPs were electrostatically coated with pectin and in this case their size increased up to 400 nm, further indicating electrostatic binding of the pectin polymer to the surface of the NPs. The surface charge (ζ-potential) of the NPs was measured to be as follows: Col IV-Ac2-26 NP (−26.33 mV ± 0.86), Col IV-Scrm NP (−24.8 mV ± 1.4), C-Col IV-Ac2-26 NP (26.6 mV ± 0.9), C-Col IV-Scrm NP (27 mV ± 0.5), P-C-Col IV-Ac2-26 NP (−25.6 mV ± 3.1), P-C-Col IV-Scrm (−23.5 mV ± 1.5), (Figure 2B). The rate of release of both the Ac2-26 and Scrm-Ac2-26 from the NPs was measured to be the fastest from the non-coated NPs, followed by the chitosan coated and the pectin-chitosan coated NPs – indicating the effect of the extra coated layers upon release of the Ac2-26 peptide (Figure 2C). The spherical shape and morphology of the NPs was confirmed with TEM, and the extra coated layers were identifiable as a darker shell layer around the NPs (Figure 2D).

**Figure 2.**
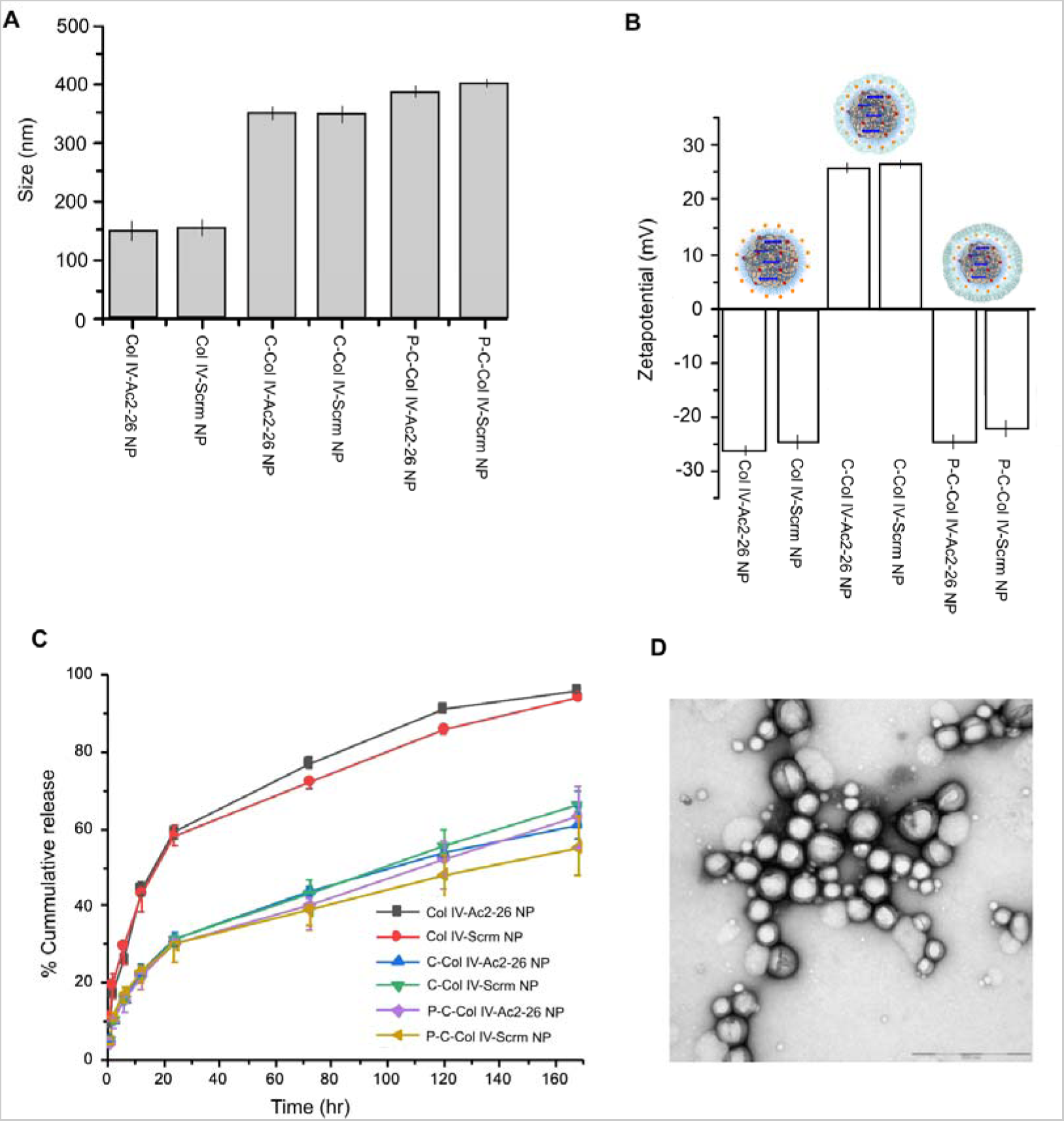
Biodegradable layer-by-layer P-C-Col IV Ac2-26 NP characterisation. Collagen IV-targeted (Col-IV) NPs encapsulating the Ac2-26 peptide (Ac2-26-NP) or controls including a randomly generated, isoelectric mismatched scrambled sequence (Scrm-NP), were developed using biodegradable polymers via a single-step nanoprecipitation method, C: chitosan, P: pectin. (A) Dynamic light-scattering of empty (NP), Ac2-26 loaded (Ac2-26-NP) and scrambled peptide (Scrm-NP) NPs were measured (mean ± SD, n = 3). (B) ζ-Potential of the NPs indicating surface charge. (C) In vitro cumulative release curve of Ac2-26 or scrambled peptide from NPs incubated at 37 °C is shown (mean ± SD, n = 3). (D) Representative TEM image of Ac2-26-NPs (scale bar = 500 nm)

### Oral administration of P-C-Col IV-Ac2-26 NP tempers postoperative colitis activity

In our murine surgical IBD model, we combined a 7 day preoperative phase of experimental DSS colitis with the surgical formation of a colon anastomosis in microsurgical technique. DSS administration led to pronounced induction of colitis, which was confirmed histopathologically in whole colonic slices (supplemental figure 1). Here, the P-C-Col IV Ac2-26-NP or P-C-Col IV Scrm-NP as control were administered through regular oral gavage every 3.5 days, starting with induction of colitis (Figure 3A). This therapeutic scheme seems appropriate, as in the clinical setting the administration of the treatment can be initiated as soon as the indication for surgery is made during acute colitis.

**Figure 3.**
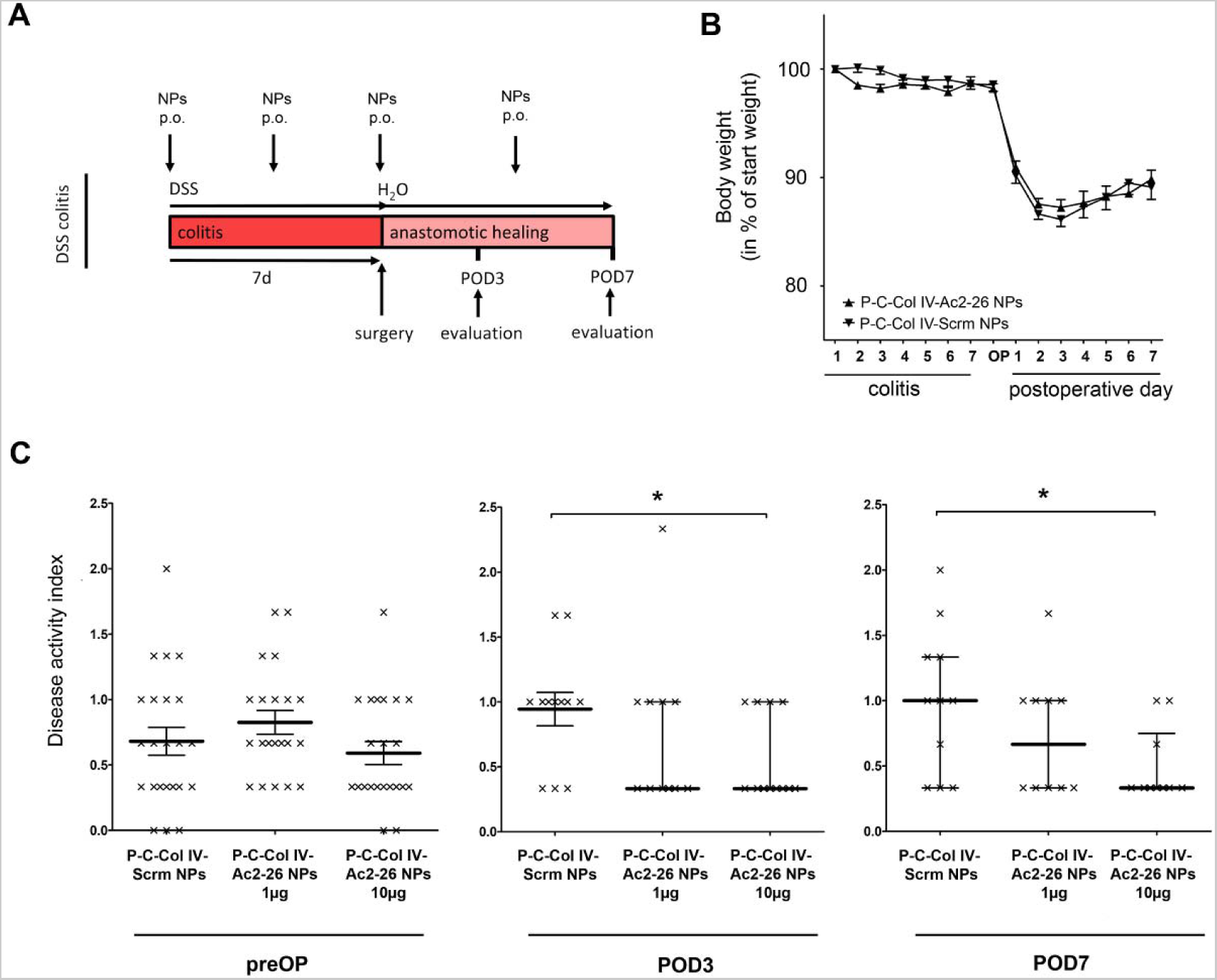
Perioperative colitis disease activity is improved by oral treatment with pectin-chitosan coated Col IV Ac2-26-nanoparticles (P-C-Col IV-Ac2-26 NPs). (A) Timeline of the experimental procedure. Mice were treated by 2 % DSS in drinking water. After 7 days DSS was changed to normal drinking water and microsurgical colonic anastomosis was performed. Mice were treated every 3.5 days with oral gavage of pectin-coated P-C-Col IV-Ac2-26 NPs in different doses (1µg Ac2-26 or 10µg Ac2-26) or pectin-coated Scrambled-nanoparticles (P-C-Col IV-Scrm NPs at a dose of 10µg of Scrm peptide) throughout the whole procedure. (B) Body weight is shown as percentage of starting weight every day during the whole procedure. (C) Disease activity index (DAI) at 3 different time points is shown: immediately preoperatively (preOP), at postoperative day 3 (POD3) and at postoperative day 7 (POD7). Statistical differences determined by Mann-Whitney-U test for nonparametric results are indicated with *(P<0.05). N=23 (P-C-Col IV-Scrm NPs), N=21 (P-C-Col IV-Ac2-26 NPs 1µg), N=22 (P-C-Col IV-Ac2-26 NPs 10µg). Three animals were excluded from analyses for reaching abort criteria (1 per each treatment group), 3 animals were excluded for intraoperative technical problems (2 in P-C-Col IV-Ac2-26 NPs 1 µg, 1 in P-C-Col IV-Ac2-26 NPs 10µg).

Mice receiving oral administrations of 10µg P-C-Col IV-Ac2-26 NP had a significantly improved disease activity index (DAI) at POD 3 (median 1 vs. 0.3, p<0.02) and POD 7 (median 1 vs. 0.3, p<0.02) compared to Scrm-NP treated mice. At the lower dose of 1µg P-C-Col IV-Ac2-26-NP there was an improvement of the DAI at POD3 and POD7, although this was not significant. Immediately before surgery, the DAI did not significantly differ between either of the treatment groups (Figure 3C). Body weight was measured as a marker of postoperative recovery, but did not differ between the treatment groups (Figure 3B).

### P-C-Col IV Ac2-26-NPs induce anastomotic wound closure and intestinal leakage

Wound closure was evaluated from the luminal side and from the abdominal side. Rigid endoscopy revealed significantly lower deciscence scores in the mice treated with 10µg Ac2-26-pNP compared to P-C-Col IV Scrm-NPs at POD 3 (median 0 vs. 2.5, p<0.02) and POD 7 (median 0.5 vs. 2, p<0.0002) (Figure 4, A-B). Scores of the mice treated with 1µg of P-C-Col IV Ac2-26-NPs were in between these groups at POD 3 and 7 demonstrating a dose dependency of the effect. Leakage from the wound causes intraabdominal inflammatory collections adjacent to the anastomosis, which lead to adhesions of the adjacent intraabdominal organs and fat tissue. Thereby adhesion formation reflects the degree of leakage during the healing process. We could observe significantly reduced adhesion scores at POD 3 (median 2 vs. 3.7, p<0.004) and POD 7 (3.7 vs. 5, p<0.02) in the mice treated with 10µg of P-C-Col IV Ac2-26-NPs compared to Scrm-NP (Figure 4C). Again, this effect was dose dependent.

**Figure 4.**
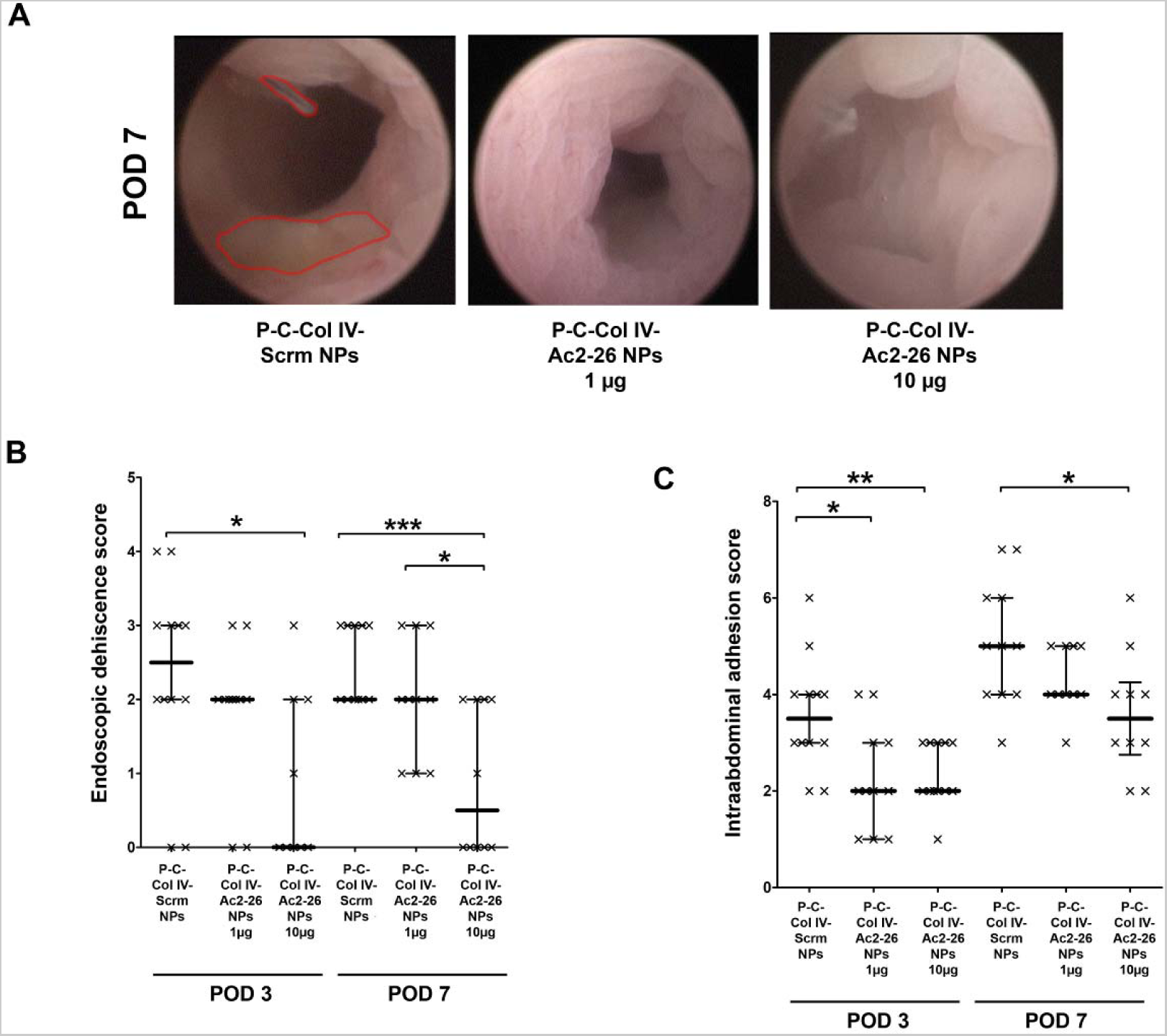
Macroscopic wound closure is improved by by oral treatment with P-C-Col IV-Ac2-26 NP. (A) Rigid colonic endoscopy was performed to analyze luminal macroscopic wound closure. Representative anastomoses at postoperative day 7 are shown. Dehiscent areas are marked in red. (B) Endoscopic deshiscence score evaluates local wound healing from a luminal perspective. Anastomotic healing was scored as described before^34^: no dehiscence = 0; sutures protruding into the lumen = 1; dehiscence less than 25% of circumference = 2; dehiscence more than 25% of circumference =3; complete dehiscence = 4. (C) Anastomotic leakage leads to adhesion formation with the adjacent tissue. Therefore grade of anastomotic leakage could be measured as follows: adhesion of mesenteric fat tissue, uterus, intestine or other organs was awarded with 1 point each. Abundance of adhesions was scored according to the following score and added: no adhesion = 0; completely bluntly removable = 1; partially bluntly removable = 2; not bluntly removable =3. Statistical differences determined by Mann Whitney U test for nonparametric results are indicated with *(P<0.05), **(P<0.01), ***(P<0.001). N=23 (P-C-Col IV-Scrm NPs), N=21 (P-C-Col IV-Ac2-26 NPs 1µg), N=22 (P-C-Col IV-Ac2-26 NPs 10µg). Three animals were excluded from analyses for reaching abort criteria (1 per each treatment group), 3 animals were excluded for intraoperative technical problems (2 in P-C-Col IV-Ac2-26 NPs 1 µg, 1 in P-C-Col IV-Ac2-26 NPs 10µg), while additionally 3 were removed from analysis in P-C-Col IV-Ac2-26 NPs 10µg at POD 3 for technical endoscopy problems.

### Orally administered P-C-Col IV-Ac2-26 NP promote the healing process in histological analysis

Additional to macroscopic evaluation of wound closure, microscopic analysis was performed to analyse the effects on the healing process at the histological level. The well-established healing score published by Phillips et al. was used for quantification of the cornerstones of regular healing: resolution of the inflammatory response, fibroblast ingrowth and subsequent collagen formation and angiogenesis.^35^ At POD 3, the healing score was higher in both doses of P-C-Col IV-Ac2-26 NP compared to the P-C-Col IV-Scrm NP control group (median 6.5 for 1µg P-C-Col IV-Ac2-26 NP, 6 for 10µg P-C-Col IV-Ac2-26 NP vs. 5.5 for P-C-Col IV-Scrm NP) (Figure 5, A-C). At POD 7, there was no significant difference between P-C-Col IV-Scrm NP treated animals and each of the P-C-Col IV-Ac2-26-NP treatment groups. These results are indicating the effect of P-C-Col IV-Ac2-26 NP especially in the early healing phase. In the functional healing score, which was adding more items including scoring the closure of the individual bowel wall layers, treatment with 10µg and 1µg P-C-Col IV-Ac2-26 NP significantly improved healing at POD 7 compared to control (Figure 5, B-C). At POD3 only the high dose of 10 µg of P-C-Col IV-Ac2-26-NP improved wound healing compared to the placebo group, but not the dose of 1µg of Ac2-26.

**Figure 5.**
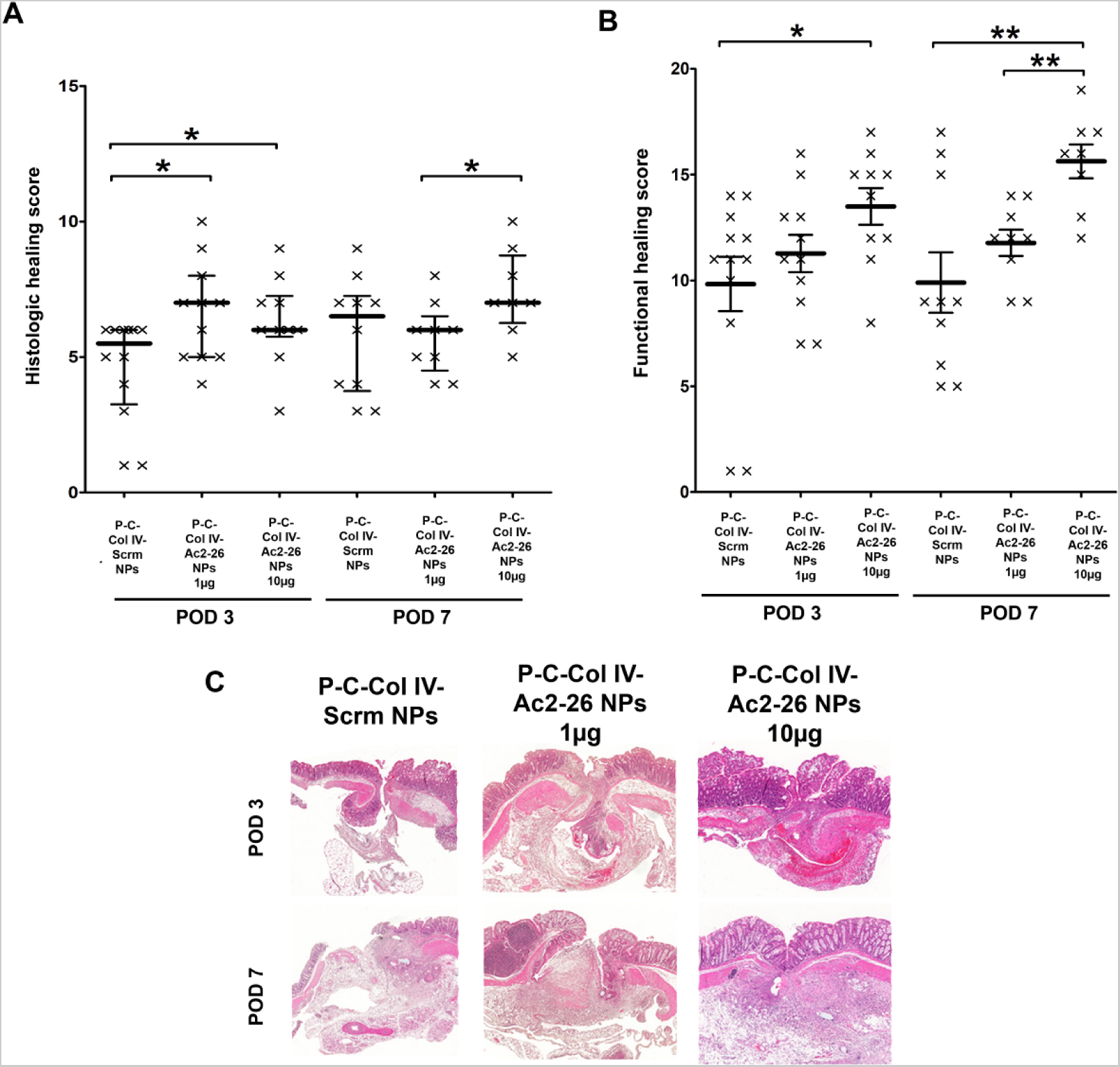
Microscopic intestinal wound healing is improved by perioperative oral administation with P-C-Col IV-Ac2-26 NPs. (A) HE-stained longitudinal sections of the anastomosis were analyzed by an experienced pathologist according to the established Phillips-Score. A score of 0 (not detectable) – 4 (highly detectable) was awarded to the following items and summed up: blood vessel ingrowth, fibroblast ingrowth, collagen formation and absence of inflammatory cells. (B) Microscopical wound closure was scored by adding additional items to the Phillips-score, focusing on closure of the individual intestinal layers^34^. (C) Representative HE-stained sections of the different groups are shown. Statistical differences determined by Mann Whitney U test for nonparametric results are indicated with *(P<0.05), **(P<0.01). N=23 (P-C-Col IV-Scrm NPs), N=21 (P-C-Col IV-Ac2-26 NPs 1µg), N=22 (P-C-Col IV-Ac2-26 NPs 10µg). Three animals were excluded from analyses for reaching abort criteria (1 per each treatment group), 3 animals were excluded for intraoperative technical problems (2 in P-C-Col IV-Ac2-26 NPs 1 µg, 1 in P-C-Col IV-Ac2-26 NPs 10µg).

## CONCLUSION

In summary, proresolving treatment with pectin coated collagen-IV-targeted Ac2-26-loaded NPs represents a novel perioperative treatment strategy with a high translational potential for patients with IBD required surgery. This approach is particularly promising additionally to established anti-inflammatory medications in the perioperative phase, as oral administration is clinically feasible, allows for gastric passage and mediates significant improvement of intestinal anastomotic healing and colitis activity.

## METHODS

### Materials and reagents

The biodegradable Poly(DL-lactic-co-glycolic acid) (50:50) (PLGA, MW: kDa), heterobifunctional NH_2_-PEG-COOH (MW: 534.6 Da) and OH-PEG-maleimide (MW: 617 Da) polymers were purchased from Nanosoft Polymers. Ac2-26 peptide (AMVSEFLKQAWFIENEEQEYVQTVK) was purchased from Tocris Biosciences. The Collagen IV targeting peptide (KLWVLPKGGGC) and the scrambled Ac2-26 peptide (WLKQKFQESVEQIAYVMENATEFEV) were purchased from Mimotopes. All NPs included a 5 molar % of PLGA.Cy5.5 fluorescent polymer for tracking studies.

### Synthesis of NPs

PLGA-PEG-COOH, PLGA-PEG-Col IV and PLGA-PEG-Cy5.5 were synthesised as previously reported.^30^ Ac2-26 or scrambled peptide loaded PLGA NPs were prepared by a single-step nanoprecipitation self-assembly method. Either Ac2-26 or scrambled peptide (120 μg) were added to the polymer mixture containing PLGA-PEG-COOH, PLGA-PEG-Col IV, PLGA-Cy5.5 (3 mg total polymer mass) in 1 mL acetonitrile, and the solution was gently vortexed overnight at RT. This polymer and peptide mixture was then added in a dropwise manner to 10mL of distilled water under gentle stirring and the solution stirred at RT for 4 hours and filtered through sterile 0.45 μm syringe filters (regenerated cellulose, 17 mm, Cole Palmer Instruments). The NPs were concentrated by centrifugation at 3000 x g for 20 min using Amicon Ultra-15 centrifugal filter units (MWCO 100KDa, Millipore Ltd), washed with deionized water, and resuspended in 1mL of H_2_O, and then further diluted with PBS prior to injection at a concentration of 1 μg Ac2-26. This solution was then dropwise added to a 1% wt ratio chitosan solution. The solution was stirred overnight and filtered through sterile 0.45 µm syringe filters (regenerated cellulose, 17 mm, Cole Palmer Instruments). The chitosan coated NPs were concentrated by centrifugation at 3000 x g for 20 min using Amicon Ultra-15 centrifugal filter units (MWCO 100KDa, Sigma-Aldrich), washed with deionized water, and resuspended in 1 mL of nuclease free H_2_O (total 3.15 mg/mL chitosan coated NP). For pectin coating the chitosan coated NPs were added dropwise to a 1 wt% solution of pectin and the NPs stirred overnight and filter the solution through sterile 0.45 µm syringe filters (regenerated cellulose, 17 mm, Cole Palmer Instruments). The pectin, chitosan coated NPs were concentrated by centrifugation at 3000 x *g* for 20 min using Amicon Ultra-15 centrifugal filter units (MWCO 100KDa, Sigma-Aldrich), washed with deionized water, and resuspended in 1 mL of nuclease free H_2_O (total 3.18 mg/mL pectin, chitosan coated NP).

### NP characterization

Ac2-26 and scrambled peptide quantification measurements were carried out on a Spark Multimode Microplate Reader. The NP mean sizes, size distribution, and z-potentials were determined by Malvern Nano ZS Zetasizer at 25 °C. All measurements were carried out in triplicates. The average particle size was expressed in intensity mean diameter and the reported value was represented as mean ± average (n=3). For TEM, a 10 μl solution of 1mg/ml freshly prepared NPs in H_2_O was deposited on carbon coated copper grids, the excess solution was blotted, and the grids were immersed in a solution of 0.75% uranyl formate stain. The stain was blotted, and the sample imaged within 1 hour of preparation on a Tecnai^TM^ G^2^ Spirit BioTwin electron microscope equipped with an AMT 2k CCD camera and low-dose software (80kV, direct mag. 98000x).

### Ac2-26 and Scrm-Ac2-26 release kinetics from NPs

To quantify the Ac2-26 release profile, 3.12 mg/mL NP samples (either P-C-Col IV-Ac2-26-NPs or P-C-Col IV-Scrm-NPs) were made in PBS, and the NPs were incubated at 37 °C. At defined time intervals, the NPs were removed, transferred to Amicon Ultra-15 centrifugal filter units (MWCO 10 kDa; Sigma-Aldrich), and centrifuged at 3,000 x g for 20 min. The NPs were then resuspended in PBS, and incubation at 37 °C was continued until the designated time point. The filtrate (10 μL) was analyzed with a nanodrop UV-Vis spectrometer, and absorbance was measured at 220 nm to determine the amount of released peptide at each time point and a cumulative % release curve generated.

### Animal experiments and study design

All animal procedures were approved by the local animal welfare committee of the regional administration of Upper Bavaria, Germany (Protocol No. 55.2-2532.Vet_02-17-203). Female BALB/c mice of 11-16 weeks with body weight between 19 and 24 g were used for the experiments (Charles River, Sulzfeld, Germany). The local animal facility environment had a temperature of 22°C and a humidity of 45-60% followed a 12-hour light/dark cycle. Animals were kept at specific-pathogen free (SPF) conditions throughout the procedure. An initial period of 1 week prior to initiation of the experiments was allowed for adaption to housing conditions.

DSS was administered for 7 days in drinking water for colitis induction which causes an inflammatory reaction by permeabilization of the mucosa for bacterial influx to the lamina propria^36^. DSS was offered ad libitum in drinking water at 2 percent (w/v) of DSS (molecular weight 36,000 - 50,000, MP Biomedicals, Heidelberg, Germany). The DAI is an established score to semi quantitatively score severity of experimental colitis by scoring weight loss, blood in stolls and stool consistency was measured prior to surgery and before sacrification.

After 7 days of DSS, the surgical anastomosis was performed and postoperatively drinking water was switched to normal water. Mice were treated throughout the whole procedure by verum ( P-C-Col IV-Ac2-26 NPs, either 1µg or 10µg) and placebo ( P-C-Col IV-Scrm-NP 10 µg) every 3.5 days throughout the whole procedure. Medication was administered by oral gavage by a 20G plastic cannula. Tissue was harvested at either POD 3 or 7 after endocopy and sacrification by isoflurane general anaesthesia and cervical dislocation.

### Surgical procedure

Anaesthesia was initiated by a mixture of 5% isoflurane (CP-Pharma, Burgdorf, Germany) and remainder of oxygen, and was maintained under reduced isoflurane concentrations of 1-3% to allow spontaneous breathing. Body temperature was maintained at 37°C by a heating pad. Subcutaneous injections of meloxicam (Boehringer Ingelheim Pharma, Ingelheim, Germany) and buprenorphine (Indivior, Berkshire, UK) were used for analgesia. After laparotomy in supine position, the colon was transected at the height of the lower renal pole, preserving the vasculature. Using an operating microscope (Carl Zeiss, Oberkochen, Germany), a standardized end-to-end anastomosis was performed in microsurgical technique by 12 single stitches of monofilamentous Ethilon 9-0 (Ethicon, Norderstedt, Germany). Postoperatively, mice had free access to regular drinking water and food *ad. libitum*. Postoperative daily assessment included body weight measurements, observation of wound healing and administration of analgesics.

### Murine colonoscopy and dehiscence scoring

Before sacrification at POD 3 or 7, colonoscopy was performedusing the Coloview system Mainz (Karl Storz, Tuttingen, Germany) with a 7 Charriere examination shaft prior to evaluation in isoflurane sedation. Instillagel (FARCO-Pharma, Cologne, Germany) was used as lubricant.The mucosa and anastomosis were evaluated under retraction of the colonoscope. Wound healing was scored according to the following score: 0 = no dehiscence, 1 = suture protruding into the lumen, 2 = dehiscence less than 25% of the circumference, 3 = dehiscence more than 25% of the circumference, 4 = complete dehiscence, as described before^34^.

### Tissue harvesting and analysis

Mice were sacrificed by cervical dislocation under isoflurane sedation. The abdomen was opened and scoring of the adhesions was performed (Range 0-7): Adhesion of mesenterial fat, the uterus, intestine or other organs was scored with 1 point each. No adhesions was scored with 0, bluntly completely removable tissue with 1, partially removable tissue with 2 and non-removable adhesions with 3. For tissue harvesting, the descending colon was resected at least 1 cm proximally and distally from the anastomosis. Prior to histological assessment, the bowel was cut in half lengthwise. Consecutive sections (4 μm thickness) of the colon bearing the anastomosis were prepared and subjected to H&E staining. Anastomotic healing was in histological sections of the anastomotic area by a pathology specialist by a healing score^35^: Scores for inflammatory cells (0-4 points, this score was inversed as higher inflammatory cell numbers reflect impaired healing), angiogenesis (0-4 points), collagen synthesis (0-4 points), fibroblast ingrowth (0-4 points) were summed up. A functional score evaluating the closure of the bowel wall was performed by adding the following items^34^: first closed layer counted from extraluminal side (0= no healed layer, 1 = serosa layer, 2 = muscular layer, 3 = submucosal layer, 4 = mucosal layer), number of healed layers (counting mucosa, submucosa, muscularis, serosa) (0 - 4), epithelial layer closed (0 = no, 1 = yes) and intact crypt architecture (0 = no, 1 = yes) were added for the functional healing score.

### Statistics

The results are presented as mean ± standard error of the mean (SEM) for parametric or median for non-parametric results. Unpaired two-tailed t-test or one-way analysis of variance (ANOVA) combined with Bonferroni post-test were applied to determine statistical significance for parametric data. For non-parametric data Mann-Whitney-U test were used. Data plotting and statistical analyses were performed in Prism (GraphPad, San Diego, USA).

## ASSOCIATED CONTENT

The Supporting Information is available free of charge at:

## ACKNOWLEDGEMENTS

This work was supported by grants of the German Research Foundation (DFG, Bonn, Germany, NE 1834/2-1 to P.A. Neumann); the Technical University of Munich (Clinician scientist scholarship, Medical School, Technical University of Munich, to S. Reischl); Imperial-TUM Collaboration Fund (Imperial College London, to N. Kamaly); TUM Global Incentive Fund (Technical University of Munich, to P.-A. Neumann).

## Notes

### Competing Interest Statement

The authors have declared no competing interest.

