## Supplementary Information for "Orally delivered biodegradable targeted inflammation resolving pectin-coated nanoparticles induce anastomotic healing post intestinal surgery"

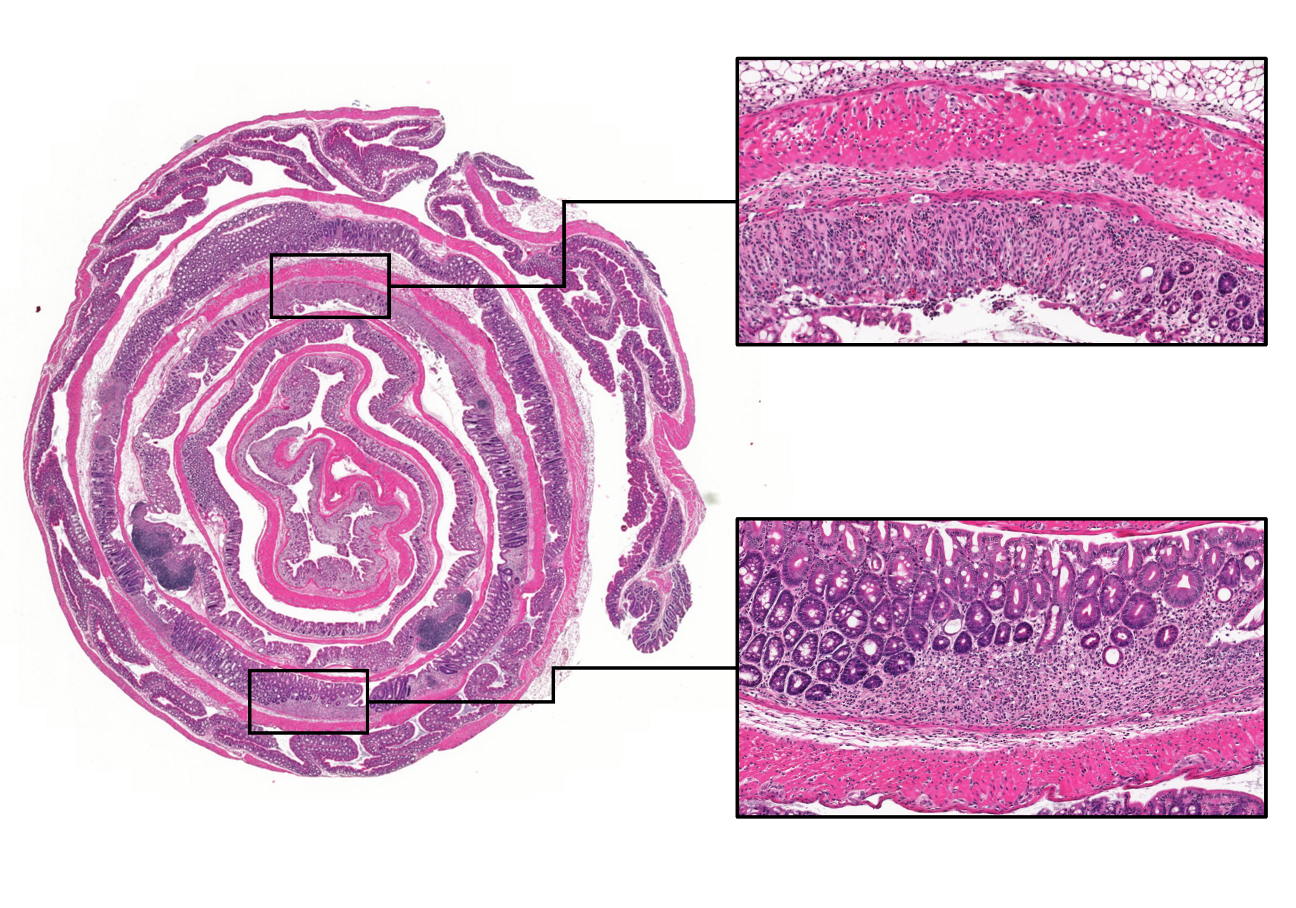

**Supplemental Figure 1. Exemplary whole colon slice after 7 days of DSS administration.** Colonic tissue was harvested, cut lengthwise, rolled, fixed and HE-stained to ensure presence of colitis. Characteristic features of DSS colitis are shown enlarged. DSS colitis is characterized by lymphocytic infiltrates, loss of crypt architecture and mucosal ulcerations.
